# Bio-Orthogonal Chemistry Enables Solid Phase Synthesis of Long RNA Oligonucleotides

**DOI:** 10.1101/2020.10.09.334060

**Authors:** Muhan He, Xunshen Wu, Phensinee Haruehanroengra, Irfan Khan, Jia Sheng, Maksim Royzen

## Abstract

Solid phase synthesis of RNA oligonucleotides which are over 100-nt in length remains to be challenging due to the complexity of purification of the target strands from the failure sequences. This work describes a non-chromatographic strategy that will enable routine solid phase synthesis of long RNA strands. The optimized five-step process is based on bio-orthogonal inverse electron demand Diels-Alder chemistry between *trans*-cyclooctene (TCO) and tetrazine (Tz) and entails solid phase synthesis of RNA on a photo-labile support. The target oligonucleotide strands are selectively tagged with Tz. After photocleavage from the solid support, the target oligonucleotide strands can be captured and purified from the failure sequences using immobilized TCO. The approach was optimized using a model 20-mer DNA strand and was successfully applied towards synthesis of 76-nt long tRNA and 101-nt long sgRNA. Purity of the isolated oligonucleotides was evaluated using gel electrophoresis and mass spectrometry, while functional fidelity of the sgRNA was confirmed using CRISPR-Cas9 experiments and flow cytometry.

## INTRODUCTION

Synthetic RNA occupies a very special place in modern research. Custom solid phase synthesis of 20-30 nucleotide-long strands has become a powerful driving force in many fields of biochemical and pharmaceutical research focusing on these relatively short RNAs. We have recently witnessed a tremendous growth of the RNA interference (RNAi) research. Scalable and high yielding solid phase syntheses of the sense and antisense siRNA strands was one of the main factors that allowed this field to mature from the discovery stage to the first-in-class FDA-approved drug, patisiran, in less than twenty years.^1–3^ In analogous fashion, facile access to short strands, containing site-specific RNA modifications, has also strongly bolstered the field of microRNA/anti-microRNA research.^4–6^

Discoveries of the twenty-first century created a strong need for a robust solid phase synthesis of longer RNA strands, over 100 nucleotides (-nt) in length. One example is non-coding RNAs, which have recently emerged as key regulators of many cellular processes. A number of important non-coding RNAs, such as human Y RNAs, are about 100-nt-long. Human Y RNAs can bind Ro60 and La proteins forming ribonucleoprotein complexes that mediate the initiation of chromosomal DNA replication.^7–8^ Synthetic non-coding RNA strands would enable investigation of the structures of the aforementioned RNA-protein complexes, the effects of RNA modifications on their stability, and mechanisms of gene regulation. Another important example, involving long synthetic RNA is Clustered Regularly-Interspaced Short Palindromic Repeats (CRISPR), one of the most effective approaches to gene editing.^9–10^ The system is composed of a 101-mer single-guide (sg) RNA that programs a nuclease (Cas9) to site-specifically cleave genomic DNA. Solid phase synthesis of sgRNA allows sequence specific incorporation of RNA modifications that can improve CRISPR efficiency, nuclease stability and reduce off-target activity.^11–12^

Despite of the strong need, solid phase synthesis of RNAs that are 100-nt in length remains to be challenging and is rarely attempted.^13–14^ The limiting step of otherwise highly optimized process is purification, illustrated in **Figure 1A**. The standard purification process entails cleavage of oligonucleotides from the solid support and concomitant deprotection of the nucleobases using AMA solution (a 1:1 aqueous solution of methylamine and concentrated ammonium hydroxide). Subsequently, desilylation of the 2-hydroxy groups is done using a fluoride reagent, such as Et_3_N·3HF. After ethanol precipitation, the target RNA strands are purified using reverse phase HPLC. The latter is the most labor intensive, time consuming and challenging step. Longer RNA oligonucleotides become increasingly more challenging to purify from failure strands that accumulate to some extent at every step of the solid phase synthesis. Long RNA strands can also be purified by preparative gel electrophoresis, which is also time-consuming and difficult to scale-up.

**Figure 1.**
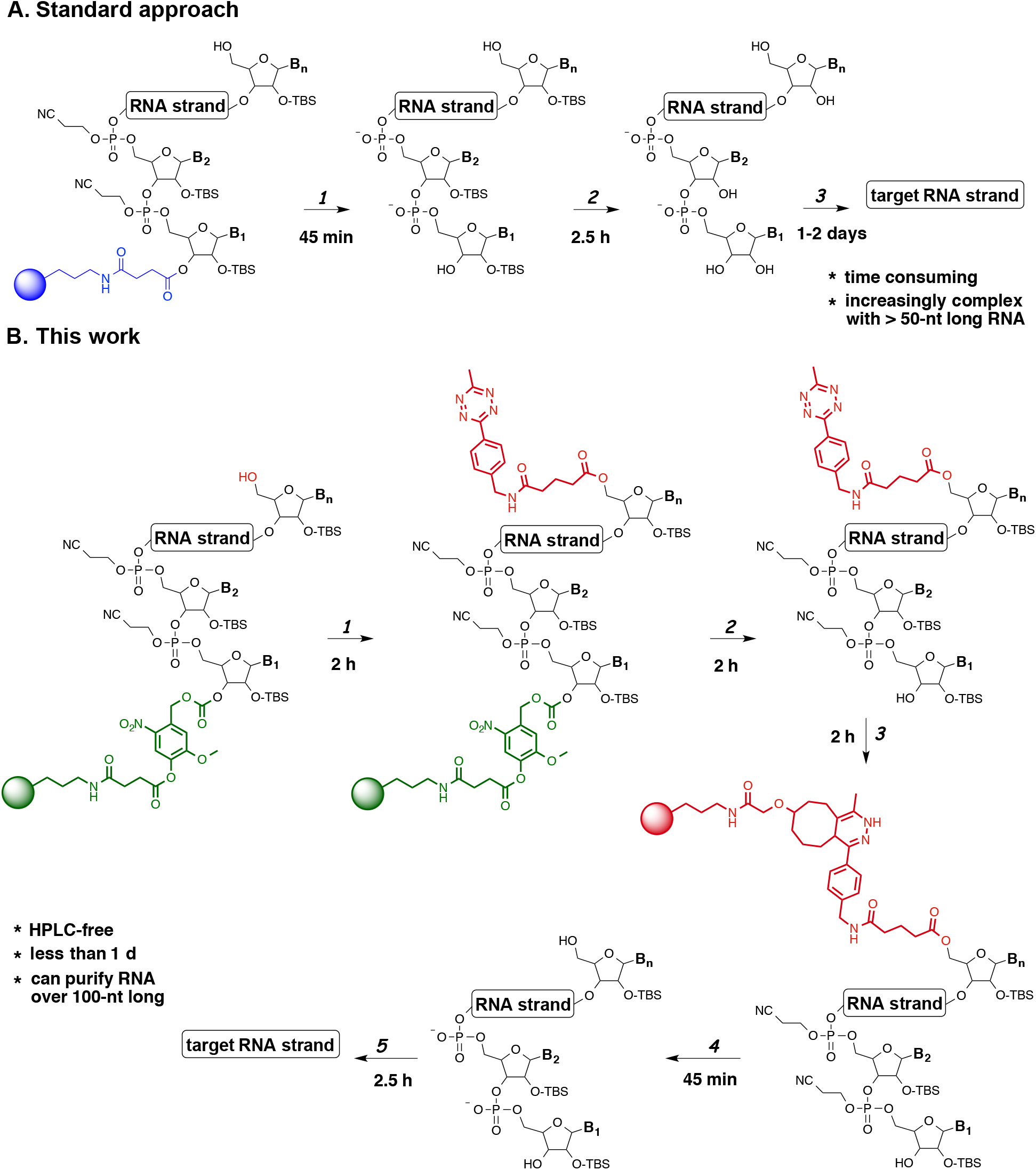
(**A**) The standard approach to isolate synthetic RNA: 1.) Cleavage and deprotection using AMA solution; 2.) Desilylation using Et_3_N·3HF; 3.) Ethanol precipitation and HPLC purification. (**B**) The non-chromatographic approach described in this work: 1.) Tagging of the target strand with Tz anhydride; 2.) Photocleavage; 3.) Capture of the target strand with CPG-TCO; 4.) Deprotection using AMA solution; 5.) Desilylation and ethanol precipitation.

A number of innovative approaches that allow HPLC-free purification of synthetic oligonucleotides have emerged. Fang described an approach for capping the failure sequences with an acrylated phosphoramidite.^15^ Subsequent polymerization of the failure sequences allows isolation of the target stand using extraction. Bergstrom reported reversible 5′-end biotinylation of synthetic RNAs.^16^ After cleavage and deprotection, the target strands were captured with NeutrAvidin coated microspheres. Beaucage synthesized DNA strands carrying 5′-siloxyl ether linkers that can be captured through an oximation reaction with aminopropylated silica gel.^17^ To our knowledge, none of these approaches have successful been applied to purify 100-nt long RNA strands that present a substantially higher degree of complexity than the ones described.

This work describes a non-chromatographic method that allows construction of long synthetic RNA strands schematically illustrated in **Figure 1B**. Our strategy is based on bio-orthogonal inverse electron demand Diels-Alder (IEDDA) chemistry between *trans*-cyclooctene (TCO) and tetrazine (Tz) that allows to selectively tag and purify structurally complex and increasing long RNA strands from the failure strands that accrue during solid phase synthesis.^18^ RNA synthesis is done on a CPG solid support modified with a photolabile linker. The linker does not impose any additional limitations on the standard solid phase RNA synthesis. After the final synthetic cycle, our approach takes advantage of the free 5’-OH group on the target strand that provides an opportunity for selective bio-orthogonal tagging. The failure strands should not have any free hydroxy groups at this stage, since they are capped during each cycle of the solid phase synthesis. Upon installing Tz on the target strand, oligonucleotides are cleaved from the solid support using UV irradiation. The target strand can be selectively immobilized using CPG-modified with TCO, while all failure strands dissolved in the supernatant can be removed. Subsequently, the target RNA strand is isolated using standard cleavage, deprotection, desilylation and ethanol precipitation steps. The 5-step process yields pure RNA strands that do not require any further HPLC purification.

## RESULTS AND DISCUSSION

Implementation the bio-orthogonal chemistry-based approach for construction of long synthetic RNA requires optimization of three key steps: 1.) immobilization of RNA on the solid support; 2.) tagging of the target RNA strands with Tz; 3.) capture of the target RNA strands. To address the first challenge, we decided to immobilize RNA on CPG using a previously reported photolabile linker. We could not utilize the standard succinate linker, as its cleavage requires AMA treatment that would inevitably also cleave Tz. Photocleavage, on the other hand, provides an orthogonal chemical procedure that preserves the tagging reagent. The photolabile linker was synthesized using procedure described by Greenberg and co-workers^19^ and attached to CPG1000, as described in **Scheme S12**.

To address the second challenge, we synthesized a series of Tz anhydrides shown in **Scheme 1A**. **Tz 1** is analogous to phenoxyacetic anhydride used commercially for ‘UltraMILD’ RNA synthesis. **Tz 2** and **Tz 3** were obtained in high yields from previously reported precursors (**Schemes S1**). Optimization of the tagging step was done using a model 20-mer DNA strand (5’-TCATTGCTGCTTAGATTGCT-3’) that was obtained in high yield using an in-house oligonucleotide synthesizer. To quantify tagging, we also synthesized a **TCO-DMT** reagent (**Scheme 1B**). Synthesis of **TCO-DMT** is described in **Scheme S2**. As illustrated in **Scheme 1B**, after the tagging step, the CPG beads were thoroughly washed to remove excess Tz and treated with **TCO-DMT** for 1h. IEDDA reaction installs DMT groups on the immobilized oligonucleotides. After removal of the supernatant, the CPG beads were treated with the detritylation reagent (3% trichloroacetic acid in CH_2_Cl_2_). Absorbance at 504 nm was measured to determine the tagging yield.

**Scheme 1.**
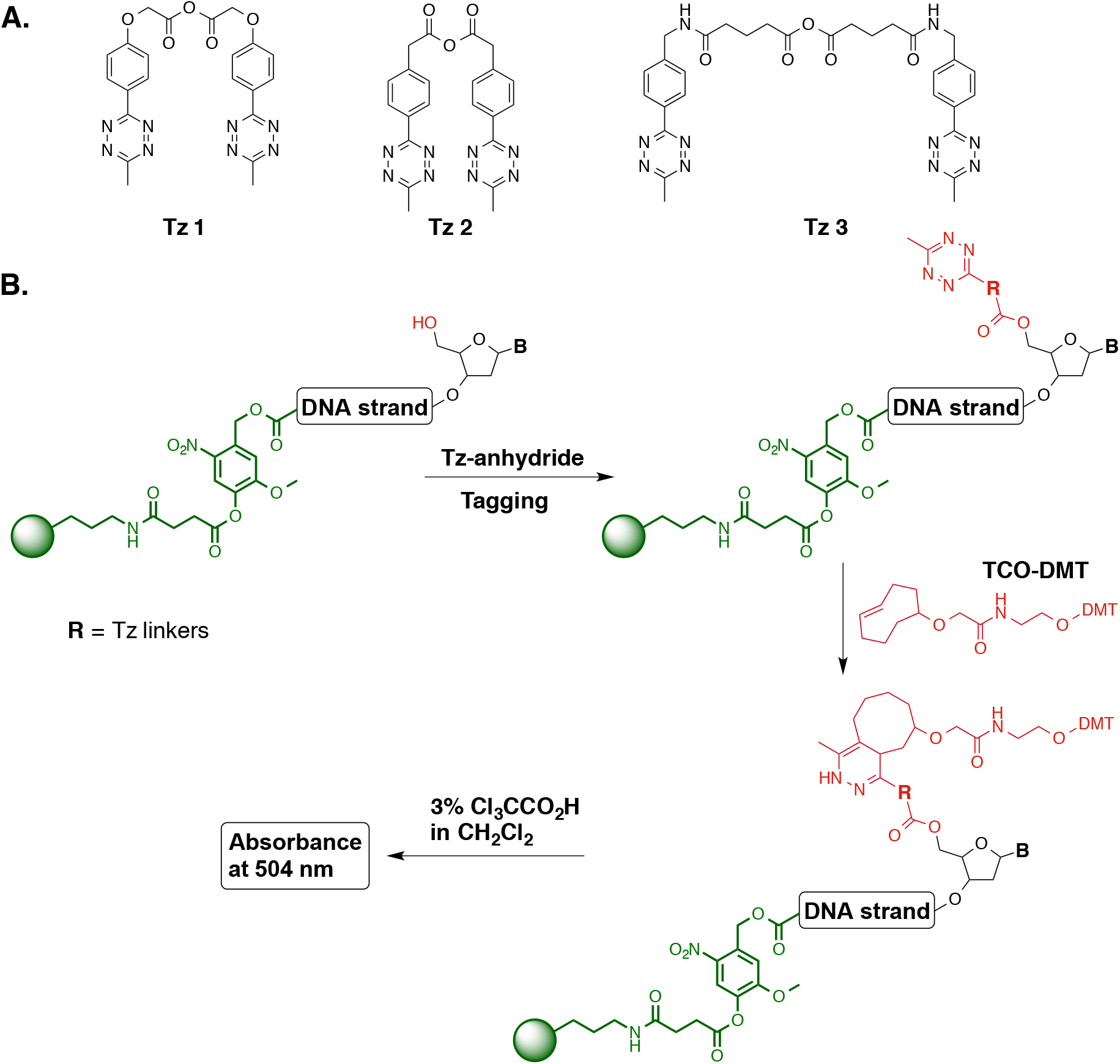
(**A**) Tz anhydride reagents that were explored for tagging of 5’-OH of oligonucleotides; (**B**) Schematic representation of the approach that was used to evaluate the efficiency of tagging.

Optimization of the oligonucleotide tagging process is outlined in **Table 1**. The model 20-mer DNA was synthesized following standard solid-phase synthesis procedure (DMT-off) with a high overall yield (>98%). We first attempted to tag the target DNA strand with **Tz 1** using standard conditions (row 1) used for capping 5’-hydroxy groups of the failure strands with acetic anhydride. Despite of considerably increasing the reaction time (the standard capping takes only 1 min), we observed a very low tagging yield. Unfortunately, alternative bases (DIPEA or DMAP) or coupling reagents did not produce any obvious improvements (rows 2-4). Therefore, we synthesized **Tz 2** and **Tz 3** to see if structural changes of the tagging reagent could provide an improvement to the tagging yield. When tagging of DNA was done in CH_2_Cl_2_ at rt using DIPEA, containing catalytic amount of DMAP, **Tz 3** significantly outperformed the other tagging reagents (rows 4-6) with a 94% yield. Increasing the temperature to 37 °C did not produce any noticeable improvement. Thus, we decided to utilize the conditions described in row 6 for the rest of the studies.

**Table 1.**
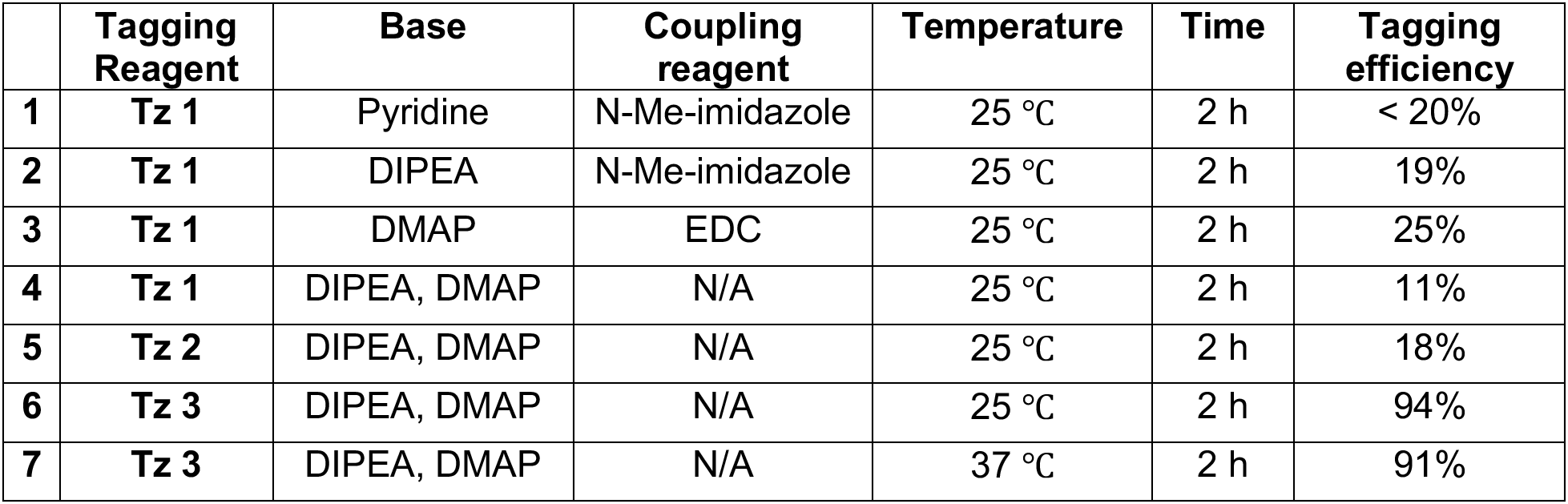
Optimization of DNA tagging using different Tz anhydrides.

To optimize the capture step, we synthesized three model DNA strands using a photolabile linker-modified CPG solid support. The 20-mer DNA strand (5’-TCATTGCTGCTTAGATTGCT-3’) containing free 5’-hydroxy group was a model target strand intended for tagging with **Tz 3**. We also synthesized two model failure strands: 17-mer DNA strand (5’-TTGCTGCTTAGATTGCT-3’) and 10-mer DNA strand (5’-TTAGATTGCT-3’), both of which were capped at the 5’-end with acetic anhydride. CPG beads containing the three model DNA strands were mixed and split into two equal portions. The first portion was processed by the standard AMA cleavage and deprotection, resulting in a crude DNA mixture (**Figure 2A**, Lane 2). The second portion was utilized for optimization of the capture process. The mixture of CPG beads was treated with **Tz 3** to selectively tag the 20-mer DNA using optimized tagging conditions. Subsequently, the beads were thoroughly washed and all three DNA strands were photocleaved using UV light. The 20-mer DNA was selectively captured by treatment with **CPG-TCO** beads. The optimal capture conditions were determined to be 2 h at 37 °C. The 17-mer and 10-mer DNA strands remained in the supernatant solution. To analyze the capture process the **CPG-TCO** beads and the supernatant solution were separated and processed by AMA deprotection. **Figure 2A** describes PAGE analysis of the experimental DNA purification process. For gel loading, each DNA sample was resuspended in water and the concentrations were adjusted based on the nanodrop measurements. Lane 2 contains a mixture of the three DNA strands (the 10-mer DNA does not stain as well as the larger strands). The successful capture of the 20-mer DNA strand was confirmed by the single dominant band observed in lane 3. Removal of the artificial failure strands in the supernatant fraction was demonstrated in lane 4. We loaded a 2-times higher concentration of DNA in lane 4 to better illustrate the removal of the failure strands (especially the difficult to visualize 10-mer DNA). The band corresponding to the 20-mer DNA strand was also observed in lane 4, indicating a partial loss of the target DNA strand during purification. We believe that this was caused by partial hydrolysis of the Tz group during photocleavage. Evidence of that has been obtained by LC-MS analysis of photocleavage products illustrated in **Figure S2**. Purification of the 20-mer DNA was further confirmed by the ESI-MS analysis shown in **Figures 2B** and **2C**. Based on the nanodrop measurements, the isolated yield of the target DNA strand was 168 nmol, which translates to 33.6% isolated yield.

**Figure 2.**
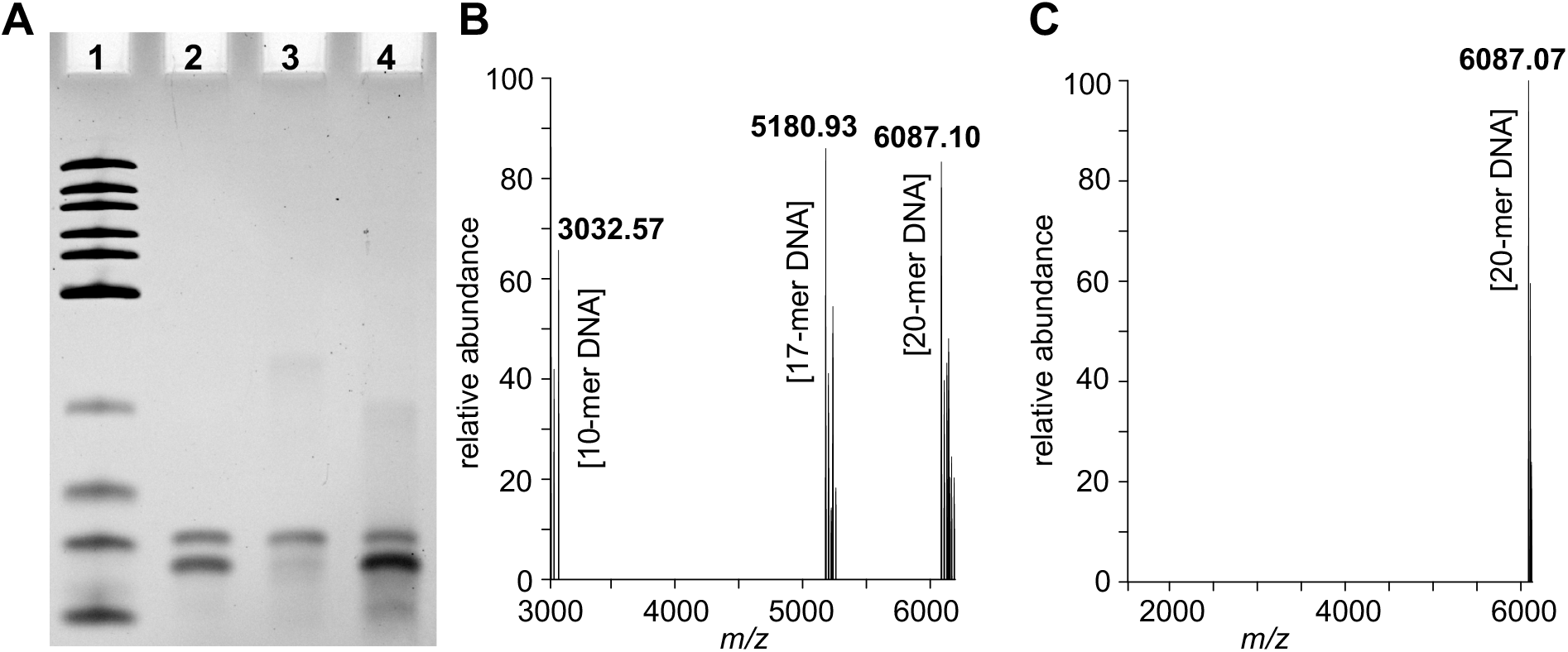
(**A**) Urea polyacrylamide gel electrophoresis analysis of capture of target 20-mer DNA. *Lane 1:* Ultra low range DNA ladder, containing DNA strands that are 300, 200, 150, 100, 75, 50, 35, 25, 20, and 10-nt long. *Lane 2:* Mixture of three model DNA strands. *Lane 3:* Captured 20-mer DNA strands. *Lane 4:* Supernatant fraction. (**B**) Deconvoluted ESI-MS spectrum of the mixture of 20-mer, 17-mer and 10-mer DNA. (**C**) Deconvoluted ESI-MS spectrum of the purified 20-mer DNA.

Once the three key steps of the non-chromatographic RNA purification have been optimized, the methodology was applied towards purification of structurally more complex RNA oligonucleotides. First, we attempted to purify a 76-nt long Lys transfer RNA (tRNA^Lys^) containing canonical nucleobases. tRNAs have a well-established, stable 3D structure. They are well known for playing important roles during protein biosynthesis, as well as its regulatory functions during numerous metabolic and cellular processes.^20^ The synthesis of tRNA^Lys^ was carried out on the photocleavable linker-modified CPG beads. The CPG beads were divided into two equal portions. The first one was processed by the standard AMA cleavage/deprotection, desilylation and ethanol precipitation, resulting in a crude mixture of target and failure RNA strands. The second portion was treated with **Tz 3** to tag the product tRNA through its 5’-hydroxy group. After tagging, RNA was cleaved from the solid support with UV light. The 76-mer tRNA was captured by CPG-TCO beads, leaving the failure strands in the supernatant. The CPG beads and the supernatant solution were separately deprotected and desilylated. After ethanol precipitation, crude tRNAs, purified tRNAs and failure strands were analyzed by urea PAGE, shown in **Figure 3**. Lane 1 contains a single stranded RNA ladder, containing strands that are 1000, 500, 300, 150, 80 and 50-nt long. The crude mixture containing the target tRNA, as well as numerous failure strands in shown in Lane 2. Purification of tRNA^Lys^ was confirmed in lane 3. Elimination of failure strands was proven by multiple fragment bands observed in lane 4. The target tRNA is also present in lane 4 due to inefficiency of the photocleavage. Based on the nanodrop measurements, the isolated yield of tRNA^Lys^ was 77 nmol, which translates to 15.4% isolated yield.

**Figure 3.**
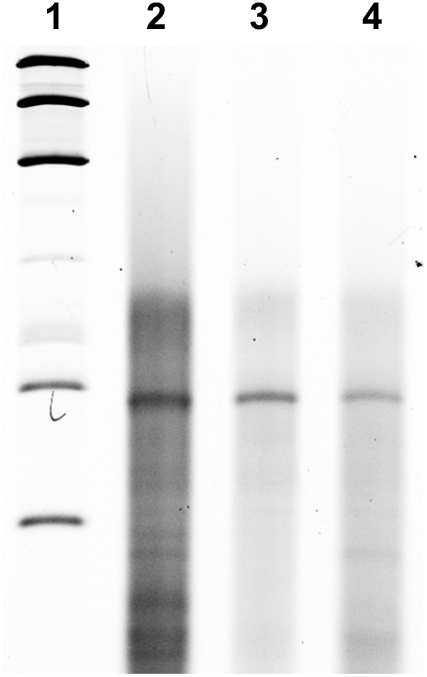
Urea polyacrylamide gel electrophoresis analysis of purification of tRNA^Lys^. *Lane 1: Low Range ssRNA Ladder*. *Lane 2:* Crude synthetic tRNA^Lys^. *Lane 3:* Purified tRNA^Lys^ strand. *Lane 4:* Failure strands.

To illustrate the practicality of our method we synthesized a 101-nt long single-guide RNA (sgRNA) frequently used for CRISPR experiments. CRISPR (Clustered Regularly Interspaced Short Palindromic Repeats) is an adaptive bacterial immune defense system comprised of a CRISPR-associated (Cas) endonuclease bound to CRISPR RNA (crRNA) cofactors that guide sequence-specific binding and subsequent phosphodiester bond cleavage of double-stranded DNA.^21^ A well-characterized and prototypical system is CRISPR-Cas9 from *Streptococcus pyogenes*.^22–23^ The guide RNA is composed of two RNAs termed CRISPR RNA (crRNA) and trans-activating tracrRNA, which can be combined in a chimeric single guide RNA (sgRNA).^21^ The 5′-end domain of sgRNA hybridizes to a target DNA sequence by Watson-Crick base pairing and guides the Cas endonuclease to cleave the target genomic DNA. The remaining double-stranded RNA structure at the 3′-end is critical for Cas9 recognition.^24^

We synthesized the sgRNA targeting GFP gene and subsequently assessed its purity and functional fidelity. The sgRNA synthesis was carried out using the photolabile linker-modified CPG solid support. After completion of the solid phase synthesis, the CPG beads were divided into two equal portions. The first one was treated with AMA for cleavage/deprotection. Followed by desilylation and ethanol precipitation, resulting in a crude mixture of target and failure RNA strands. The second portion was purified using the HPLC-free process described above. Purification was characterized by 15% urea PAGE, shown in **Figure 4**. Lane 1 contains a single stranded RNA ladder, containing strands that are 1000, 500, 300, 150, 80 and 50-nt long. Lane 2 represents the crude sgRNA containing many impurity bands. The purified sgRNA shown in lane 3. The removed failure sequences are in lane 4. Based on the nanodrop measurements, the isolated yield of sgRNA was 50 nmol, which translates to 10% isolated yield.

**Figure 4.**
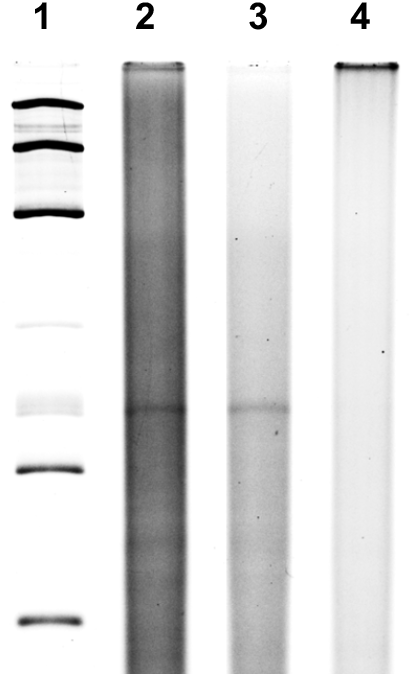
Urea polyacrylamide gel electrophoresis analysis of purification of sgRNAs. *Lane 1: Low Range ssRNA Ladder*. *Lane 2:* Crude synthetic sgRNA. *Lane 3:* Purified sgRNAs. *Lane 4:* Failure strands.

To confirm the functional fidelity of the purified sgRNA we carried out CRISPR-Cas9 experiments targeting the GFP gene in HEK293T cells, expressing GFP and Cas9.^25^ The cells were transfected with the purified sgRNA that would guide Cas9 to produce double-stranded DNA breaks at the GFP site. Thus, a decrease of GFP expression level would be expected if the purified sgRNA is properly functioning. As positive control, the cells were transfected with purchased HPLC-purified crRNA and tracrRNA that form a two-component structure that has been reported to function analogous to sgRNA.^25^ After the three-day transfection, the cells were fixed and GFP expression was evaluated using flow cytometry, shown in **Figure 5**. Prior to the transfections, 65% of HEK293T cells were determined to express GFP (*purple curve*). The percentage of cells expressing GFP decreased to 42% after transfection with the commercial crRNA and tracrRNA mixture (*red curve*). Transfection of HEK293T cells with sgRNA purified by our experimental procedure resulted in a comparable attenuation of GFP expression, as illustrated in **Figure 5**. In the latter case 49% of HEK293T cells were determined to express GPF (*green curve*). These results indicate that our experimental RNA purification procedure achieved sgRNA with sufficient functional integrity.

**Figure 5.**
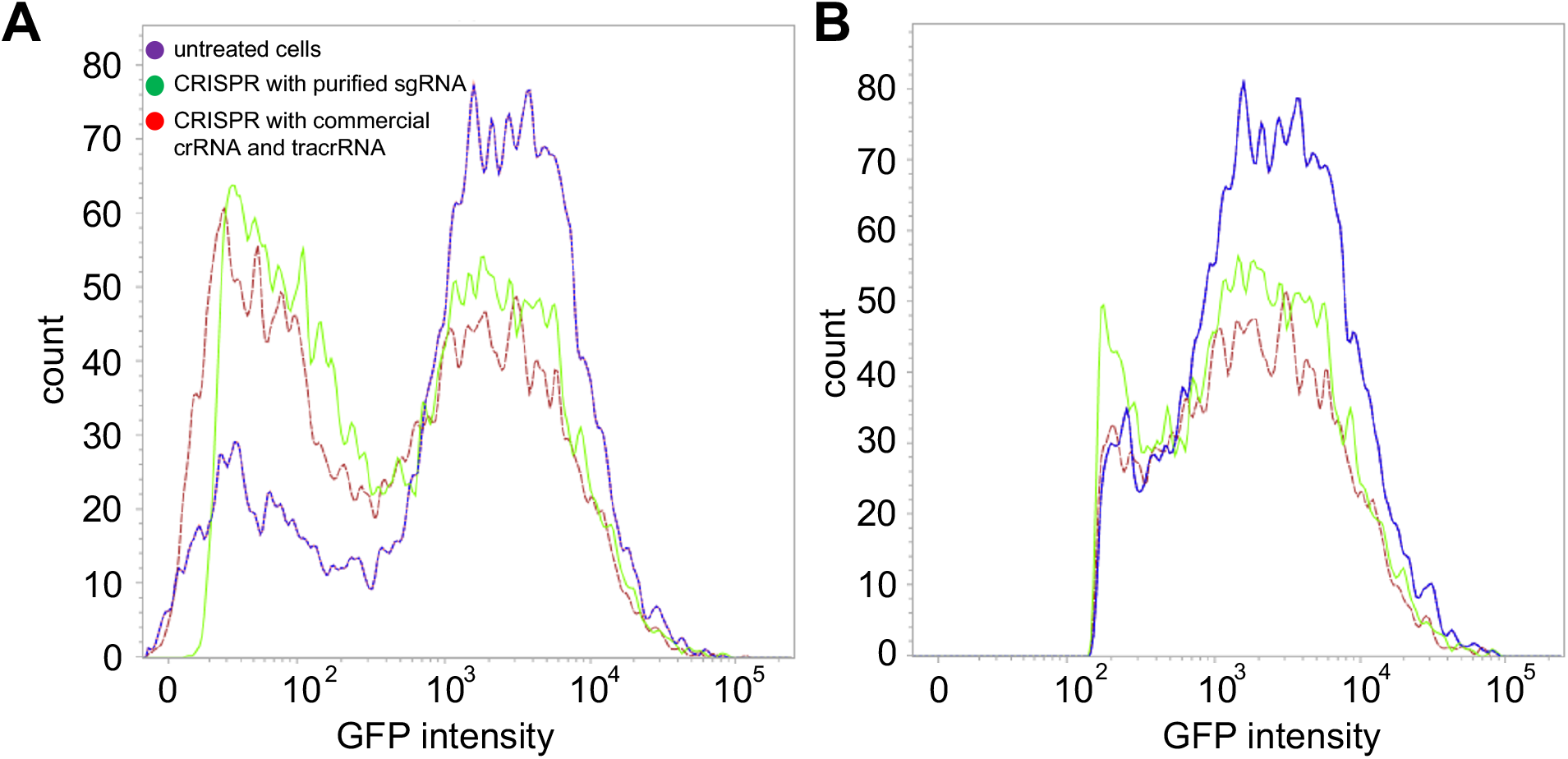
Flow cytometry analysis of CRISPR experiments targeting the GFP gene in HEK293T cells, expressing Cas9. (**A**) Histograms of total GFP-expressing cells: untreated cells are shown in purple, cells transfected with the commercial HPLC-purified crRNA and tracrRNA are shown in red, cells transfected with the sgRNA purified using our experimental procedure are shown in green. (**B**) Histograms of total live cells using the same coloring scheme.

## CONCLUSION

This report describes development of the methodology that allows construction of RNA strands that are over 100-nt in length. Our ultimate goal is to convince the RNA community that our approach can expand the scope of solid phase RNA synthesis towards long and more complex oligonucleotides. Our methodology is based on the bio-orthogonal reaction between *trans*-cyclooctene and tetrazine that are highly selective for each other and have minimal cross-reactivity with other functional groups found in RNA. As the result, lEDDA chemistry does not interfere with the intricate structural elements of RNA. The bio-orthogonal click chemistry is highly efficient, even at very low concentrations of TCO and Tz. Thus, it can still be effectively applied towards isolation of long synthetic RNA strands obtained in low overall yield. The key steps of our procedure were optimized using model DNA strands. The optimized protocol was implemented towards purification of the 76-nt long tRNA and 101-nt long sgRNA. In addition to determining purity, the functional fidelity of the 101-nt long sgRNA was characterized using CRISPR experiments. While optimizing our procedure, we discovered a critical drawback associated with a partial loss of the Tz-tagged oligonucleotides during photocleavage. This drawback lowers the yield of the isolated target RNA. In our future work, we plan to address this by designing new photolabile linkers that can be cleaved using a more chemically benign longer-wavelength light.

## ACKNOWLEDGMENTS

This work was supported by the NIH grant R21CA228997-01 to M.R. and the NSF grant 1664577 to M.R. The authors would like to thank Prof. Greenberg (Johns Hopkins University) for his suggestions regarding synthesis of the photolabile linker. The authors would also like to thank the staff of the Life Sciences Research Building at the University at Albany for their help in acquiring the experimental data.

